# The Nox2 NADPH oxidase regulates neutrophilic inflammation in the oral cavity

**DOI:** 10.1101/2025.08.26.671820

**Authors:** Shunying Jin, Richa Singhal, Jianzhu Luo, Kelley N Cooper, Marina Terekhova, Katherine A Carey, Maxim N Artyomov, Venkatakrishna R Jala, Rachel A Idol, Mary C Dinauer, Richard J Lamont, Juhi Bagaitkar

**Affiliations:** Department of Oral Immunology and Infectious Diseases, University of Louisville, Louisville, Kentucky; Center for Microbe and Immunity Research, Abigail Wexner Research Institute, Nationwide Children’s Hospital, Columbus, Ohio; Department of Pathology and Immunology, Washington University School of Medicine, St. Louis, Missouri; Department of Microbiology and Immunology, School of Medicine, University of Louisville, Louisville, Kentucky; Department of Pediatrics, Washington University School of Medicine, St. Louis, Missouri; Department of Pediatrics, College of Medicine, The Ohio State University, Columbus, Ohio

**Author notes:** **CORRESPONDING AUTHOR: Juhi Bagaitkar, PhD,** 700 Children’s Drive, WA 4.111, Columbus, OH 43205. Co-first authors (Shunying Jin, Richa Singhal).

## Abstract

The leukocyte NADPH oxidase 2 (NOX2) is an important regulator of inflammatory responses, independent of its antimicrobial activity. Inactivating mutations in NOX2 cause chronic granulomatous disease (CGD), a severe immunodeficiency associated with recurrent infections and dysregulated neutrophilic inflammation. Recurrent oral ulcers, stomatitis, gingivitis, and other inflammatory issues affecting the oral mucosa have been observed in patients with CGD; however, the underlying mechanisms are not known. Here, we present evidence that the extensive inflammatory destruction of oral mucosal tissues observed in Nox2-deficient or *Cybb^KO^*mice was not caused by impaired antimicrobial surveillance against oral pathobionts but instead resulted from a cell-intrinsic dysregulation of neutrophil inflammatory responses. Transcriptional and cellular profiling of oral tissues isolated from wild-type and *Cybb^KO^* mice showed a dominant neutrophil signature, which was accompanied by a significant upregulation of several bone-resorbing, tissue-degrading inflammatory cytokines and a reduced expression of nuclear factor erythroid 2-related factor 2 (Nrf2) regulated genes. Mechanistically, hyperinflammatory responses were mitigated by restoring Nrf2 transcriptional activity using a synthetic agonist. Thus, our studies show that Nox2 oxidase and its derived reactive oxygen species are crucial for balanced recruitment and cell-intrinsic regulation of neutrophil inflammatory responses within oral tissues in an Nrf2-dependent manner.

## INTRODUCTION

The leukocyte NADPH oxidase 2 (Nox2) is a multi-subunit enzyme complex predominantly expressed in phagocytes. The activation and assembly of the Nox2 complex on plasma or phagosomal membranes generate superoxide (O_2_^−^), a precursor to antimicrobial reactive oxygen species (ROS) that have critical immunoregulatory roles. Inactivating mutations in the Nox2 oxidase subunit genes are associated with chronic granulomatous disease (CGD), a severe immunodeficiency characterized by life-threatening infections and an excessive inflammatory response ^1,2^. Dysregulated inflammatory responses are a significant aspect of CGD pathology, and patients frequently present with sterile granulomatous inflammation in the lungs, gastrointestinal tract, and mucocutaneous lesions. Interestingly, patients with CGD are also prone to oral inflammation characterized by stomatitis and recurrent oral ulceration affecting the cheeks, lips, the floor of the mouth ^3–5^, and gingival inflammation ^6^. Despite the prevalence of oral mucosal inflammation in patients with CGD, insights into the underlying mechanisms are lacking.

The Nox2 is highly expressed in neutrophils, a highly abundant cell type in oral mucosal tissues. They are actively and continuously recruited into the oral cavity, a hostile environment rich in diverse activating stimuli such as bacterial and fungal ligands, food antigens, allergens, and ongoing tissue damage due to mastication. Within the gingival tissues, neutrophils limit microbial numbers and barrier breaches by phagocytosing bacteria, degranulating, and forming neutrophil extracellular traps (NETs), thus playing an important role in antimicrobial surveillance. Their presence is also essential for wound healing and immune homeostasis upon efferocytic clearance ^7,8^. However, the cell-intrinsic mechanisms that restrain neutrophil effector responses in the oral cavity to prevent tissue damage via excessive activation under homeostatic conditions remain unknown. A substantial body of literature offers compelling evidence that Nox2-derived ROS, independent of their antimicrobial role, play a vital role in modulating neutrophil inflammatory potential within tissues ^9^. Interestingly, neutrophils lacking Nox2 exhibit a primed or activated phenotype with enhanced azurophilic granule exocytosis, which is coupled with the excessive release of tissue-degrading proteases ^10^. The absence of Nox2 activity was also associated with elevated calcium entry in neutrophils and increased levels of the inflammatory lipid mediator leukotriene B4 ^11,12^. Other studies demonstrate that excessive activation of the transcription factor NF-κB (Nuclear factor-kappa B) led to overproduction of many pro-inflammatory cytokines, higher inflammasome activity ^13,14^ that correlated with increased IL-1 production in neutrophils and monocytic cells ^15–17^. We previously showed that Nox2 deficiency was also linked to the excessive mobilization of neutrophils from bone marrow pools, resulting in their increased accumulation within the inflamed tissues of Cybb/Nox2 knockout (*Cybb^KO^*) mice, resulting in a significant delay in resolving the inflammatory response^16–20^.

Here, we examined whether Nox2 and its derivative ROS are crucial in controlling inflammatory responses at the oral mucosa, a barrier surface heavily populated by microbes. Our mechanistic studies in mouse models, as reported below, show significant inflammation characterized by erosion of oral soft tissues and the recession of alveolar bone, which occurs in *Cybb^KO,^* is primarily driven by the overabundant recruitment of neutrophils into oral tissue and hyperactivation. Interestingly, despite the presence of a dysbiotic microbial community, we found no signs of sepsis or dissemination of periodontal pathobionts systemically. Thus, our findings highlight a previously uncharacterized role for Nox2 in cell-intrinsic regulation of neutrophil responses within oral tissue, an environment replete with varied activating stimuli.

## METHODS

### Mice

We purchased C57BL/6J wild-type (WT) mice from the Jackson Laboratory. Dr. Mary Dinauer provided Cybb/Nox2 knockout (*Cybb^KO^*) mice ^21^ and conditional knockout mice lacking Nox2 activity in neutrophils (*Cybb*^f/f^ *Ncf2*^f/f^) ^22^. Dr. Krishna Jala provided Nrf2 knockout (*Nfe2l2^−/−^*) mice. Both male and female mice, aged 6 to 10 weeks, were used for all studies to account for sex as a biological variable. The Institutional Animal Care and Use Committees (IACUC) at the University of Louisville and the Abigail Wexner Research Institute at Nationwide Children’s Hospital approved all experiments.

### Ligature model of oral inflammation

Gingival inflammation was induced by placing a 5-0 silk ligature around the second maxillary molar of mice ^23^. After 8 days, the mice were euthanized, and the gingival tissues surrounding the ligatures were isolated and either flash-frozen for RNA extraction or enzymatically digested for flow cytometry ^24^. Mouse heads were fixed in 10% formalin and scanned using microcomputed tomography (µCT). Average bone loss was measured by taking linear measurements (in millimeters) from the cementoenamel junction (CEJ) to the alveolar bone crest (ABC) in the interdental regions between the first and second molars (M1-M2) or the second and third molars (M2-M2’) (**Figure S1A-B**) and represented as CEJ-ABC length, as Park et al. ^25^ previously described. In specific experiments, mice received 25 mg/kg sulforaphane (SFN) intraperitoneally (i.p.) once daily for seven days after placing ligatures ^26^. For experiments related to microbiome analysis, female WT and *Cybb^KO^* mice were co-housed for a month prior to ligature placement.

### Bacterial peritonitis

Mice were challenged with either 10^7^ colony-forming units (CFUs) of *P. gingivalis* 33277 or 10^7^ CFU of ligature-associated oral bacteria i.p.) and euthanized after 4 h. Peritoneal cavities were lavaged with 4 ml sterile PBS + 2 mM ethylenediaminetetraacetic acid (EDTA). Lavage fluid (50 µL) was plated on trypticase soy broth (TSB) blood agar plates to estimate bacterial CFUs (in vivo survival). Separately, lavage fluid was analyzed for inflammatory cytokines by 32-plex cytokine arrays, and leukocyte differentials were determined by flow cytometry.

### Bacteria

*Porphyromonas gingivalis* 33277 was cultured anaerobically in enriched trypticase soy broth (eTSB) supplemented with hemin (5 μg/mL), menadione (1 μg/mL), and 1 mg/mL yeast extract. *Fusobacterium nucleatum* 25866 was cultured anaerobically in brain-heart infusion broth (BHI) with yeast extract (1 mg/ mL), hemin (5 μg/mL), and menadione (1 μg/mL). *Filifactor alocis* was cultured anaerobically in BHI with yeast extract (1 mg/mL), hemin (5 μg/mL), menadione (1 μg/mL), cysteine (0.05%), and arginine (0.05%). All strains were grown at 37 °C with 85% N_2_, 10% H_2,_ and 5% CO_2_. Ligatures were isolated from WT and *Cybb*^KO^ mice, placed in 1 mL of sterile phosphate-buffered saline (PBS), and vortexed to dissociate ligature-associated bacteria. Suspension (0.5 mL) was added to two tubes containing 10 mL of eTSB to cultivate ligature-associated bacteria and then incubated either aerobically or anaerobically overnight. The next day, 10^7^ CFU of aerobically and 10^7^ CFU anaerobically cultured bacteria were combined in PBS and injected into mice i.p.

### Degranulation assays

Degranulation of mouse bone marrow neutrophils (BMNs) was measured by assessing the mobilization of granule markers to the plasma membrane by flow cytometry. BMNs were challenged with various bacteria at a multiplicity of infection of 1:10 and 1:100 (cell: bacteria) at the indicated time points. At the end of each time point, cells were washed twice with 3 mL of ice-cold flow buffer (1% bovine serum albumin, 2 mM EDTA in PBS). For flow staining, cells were resuspended in 50 μL of 2.4 G2 supernatants to block Fc receptors for 10 mins at room temperature and then stained with the fluorochrome conjugate anti-mouse Ly6G-V450, CD63-PE, and CD35-BV605 antibodies for 1 hour on ice. Data were collected on FACS Celesta (BD Biosciences) and analyzed with FlowJo V10.

### Data availability

RNA sequencing data have been submitted to the Gene Expression Omnibus database under Accession No. GSE184556. The metagenomics data are in SRA PRJNA1235937.

### Statistical analysis

GraphPad Prism, Version 9.3.1, was used to perform all statistical analyses. A detailed description of the specific statistical test used for data analysis, as well as post-hoc correction, is provided in the legend to each figure.

**Please see Supplemental Materials for additional methods.

## RESULTS

### Nox2 deficiency enhances neutrophil recruitment and inflammatory damage in oral tissues

To reproducibly model oral inflammation in mice, we used the well-characterized ligature-induced periodontitis (LIP) ^23^. In this model, ligature placement around the second maxillary molars leads to microbial accumulation, immune cell infiltration, and localized bone resorption (**Figure 1A)**. We observed a significantly higher mobilization of neutrophils into the blood within 24 h of induced oral inflammation (ligature placement) in Nox2/Cybb knockout (*Cybb*^KO^) mice ^21^, which tracked with significantly higher neutrophil accrual in the gingival tissues (**Figure 1 B-F**). To determine the inflammatory mediators in serum, we performed a multiplexed 32 cytokine assay and observed an early spike in factors linked with neutrophil mobilization, such as granulocyte colony-stimulating factor and CXCL1 (KC) ^27^ that remained high until Day 8 in the blood of *Cybb*^KO^ mice, but not WT mice (**Figure 1G-H**). This was consistent with our previous observations in other tissues, such as sterile peritonitis ^16^ or zymosan-induced lung injury ^17^, where the absence of Nox2 activity was associated with an unbalanced neutrophil response to experimentally induced inflammation similar to that observed in LIP. Excessive neutrophil recruitment and their prolonged activation contribute significantly to localized inflammatory bone loss in the oral cavity ^28^. Dysregulated neutrophil responses not only lead to soft tissue ulceration but are also associated with inflammatory bone recession in mice ^23,29^ as well as patients with neutrophil immunodeficiencies ^30^. µCT analysis of mouse skulls on Day 8 demonstrated pathological resorption of alveolar bone around the ligated molars, which was significantly more severe in *Cybb*^KO^ mice (**Figure 1I-K**). Average bone loss was measured by taking linear measurements (in millimeters) from the CEJ to the ABC in the interdental regions around the ligature, between the first and second molars (M1-M2) and the second and third molars (M2-M2’) as illustrated and quantified in **Figure S1 A-B**.

**Figure 1:**
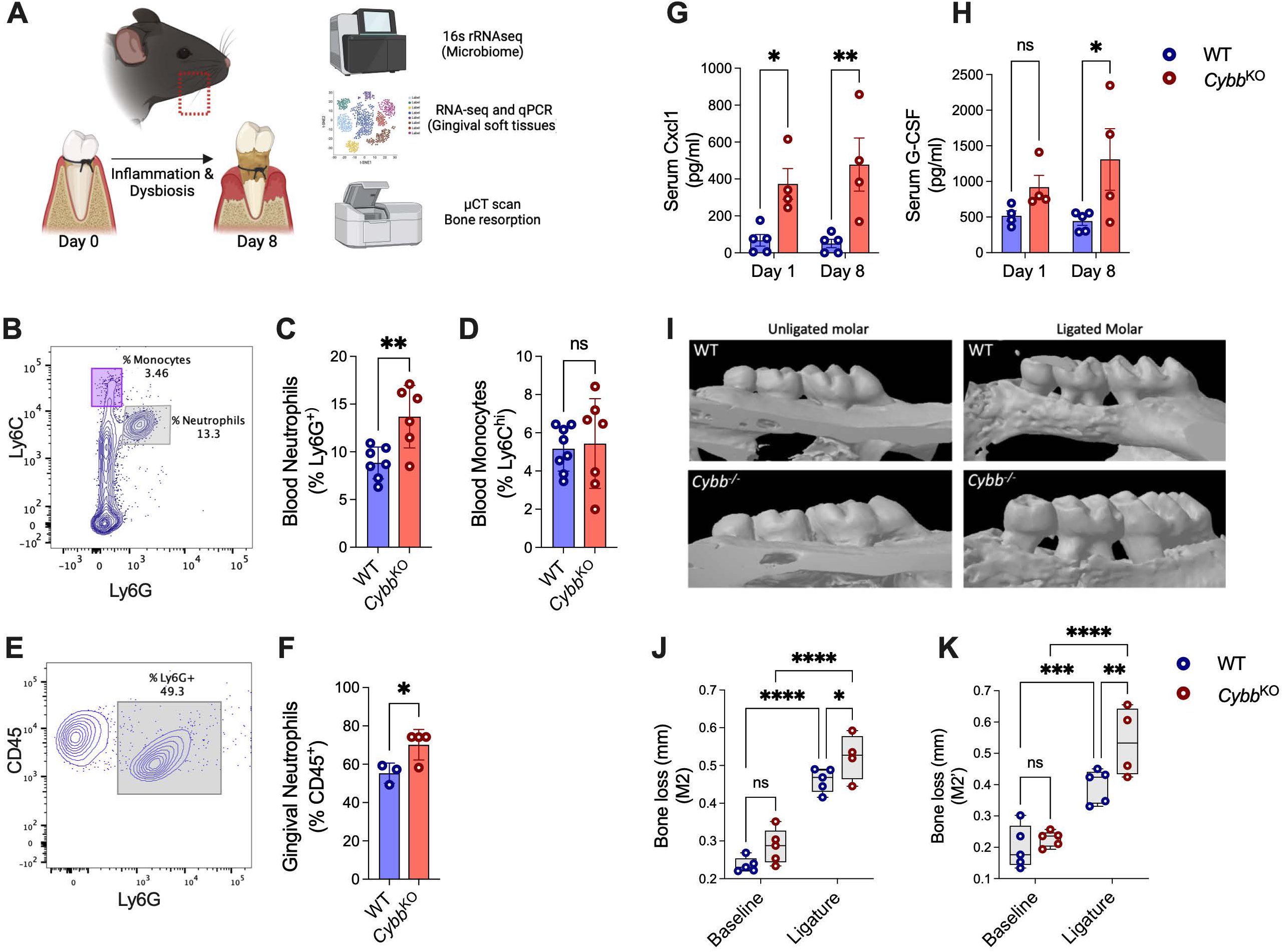
Nox2 deficiency drives excessive neutrophil recruitment and damage in oral mucosal tissues. **(A)** Illustration depicting the ligature-induced periodontitis in mice on the left second maxillary molars of wild-type (**WT**) and *Cybb^KO^* mice. (**B-D**) 24 h after ligature placement, the relative abundance of neutrophils (Ly6G^hi^ Ly6C^int^) and monocytes (Ly6C^hi^ Ly6G^−^) were determined in peripheral blood via flow cytometry. (**E-F**) Gingival neutrophil numbers on Day 8. Data from 4-7 mice per group are displayed as mean ± SD, and a t-test determined statistical differences (*P<0.05; **P<0.01). (**G-H**) Multiplexed 32-cytokine arrays determined serum levels of KC/ CXCL1 and granulocyte colony-stimulating factor (G-CSF) on Day 1 and Day 8 post-ligature placement. (**I-K**) Average bone loss (in millimeters) in the interdental regions (M2, M2’) of the ligated molar and unligated contralateral control molars from the same mice. Data from 5-8 mice per group are displayed as mean ± SD, and statistical differences were determined using two-way ANOVA with Sidak’s multiple comparison tests (*P<0.05; **P<0.01; ***P<0.001; ****P<0.0001).

To further delve into the underlying dysregulated molecular pathways, we performed RNA-seq of the gingival tissues. Volcano plots compared the variation in differentially expressed genes (**DEGs**) in ligated or inflamed gingival tissues (**Figure 2A**) and gingival tissues from naïve WT and *Cybb^KO^*mice (**Figure S1C**). Interestingly, no significant changes were noted in gene expression in the oral tissues of naïve WT vs. *Cybb^KO^* mice (**Figure S1C**). Furthermore, we did not observe any spontaneous bone loss in the unligated/naive *Cybb^KO^* mice, indicating that ROS deficiency does not predispose to infections from oral bacteria (**Figure 1I-K**). In contrast, the gingival tissues of ligated/inflamed mice indicated significant inflammatory gene signatures with many more DEGs and a strong inflammatory response in Nox2-deficient animals (**Figure 2A**). We performed gene-set enrichment analysis in the order of decreasing log2 fold change in a ranked list of all genes. Gene ontology (GO) analysis indicated that the most significantly enriched terms *(according to q values or NES)* were ‘neutrophil degranulation’ and ‘signaling by interleukins’ (**Figure 2 B-D**). The top ten enriched pathways are presented in **Figure 2B**.

**Figure 2:**
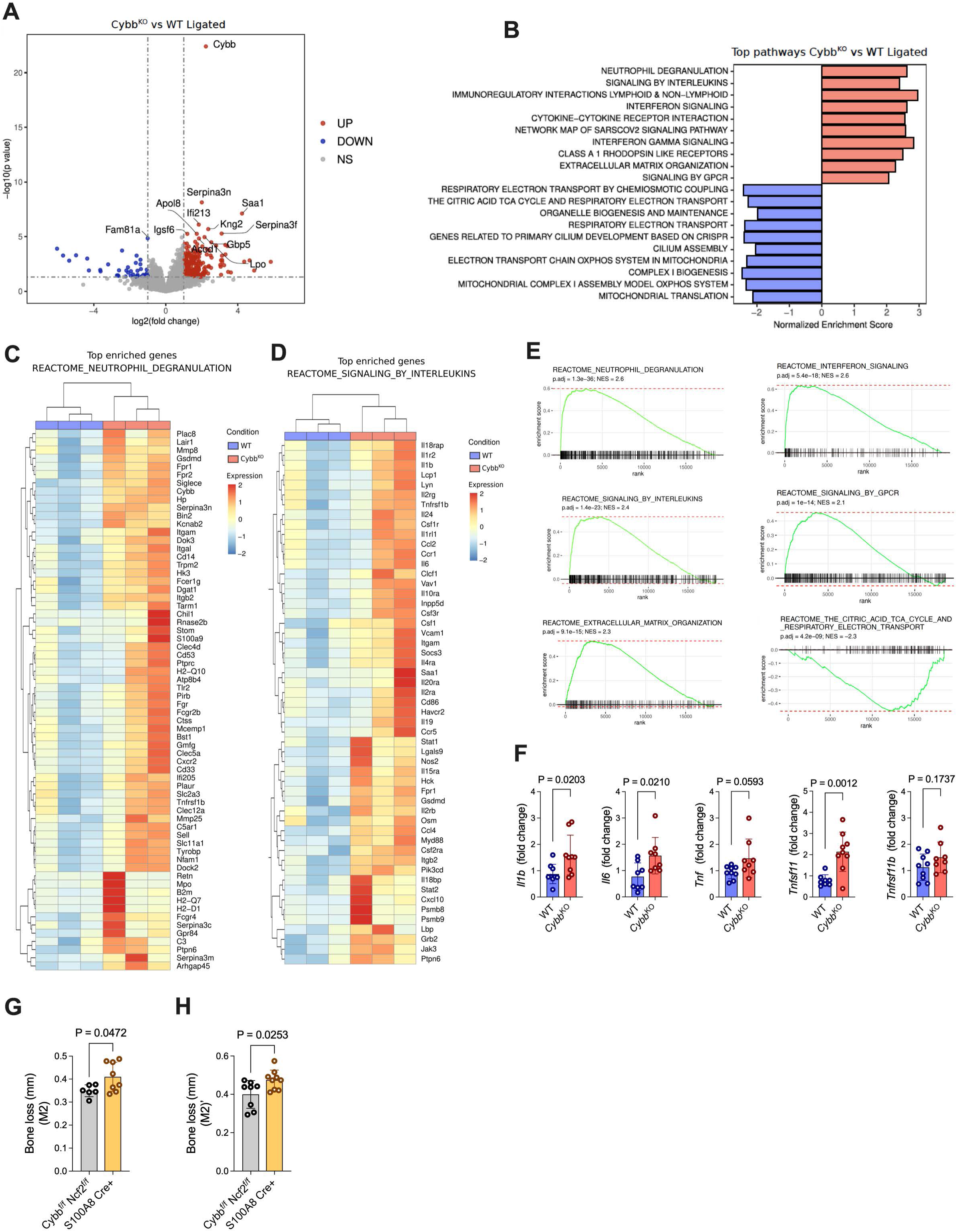
Hyperinflammatory gene signatures predominate in inflamed gingival tissues of Nox2-deficient (*Cybb^KO^*) mice. Gingival tissues of wild-type (**WT**) and *Cybb^KO^* mice were extracted on Day 8 and analyzed by RNA-seq. (**A**) Plotting log2 (fold change) versus –log10 (adjusted p-value) generated volcano plots of RNA-seq data, representing differences between ligated WT and *Cybb^KO^*. **(B)** Differentially expressed genes (**DEGs**) were used to determine enriched and suppressed pathways. Heatmaps of top DEGs in the (**C**) Reactome_Neutrophil_Degranulation and (**D**) Reactome_Signaling_By_Interleukins. (**E**) Gene set enrichment analysis (**GSEA**) plots of the major enriched pathways. (**F**) Validation of select genes by **qPCR** from gingival tissues. Data from 4-7 mice per group are displayed as mean ± SD, and statistical differences were determined using t-test. (**G-H**) Average bone loss (in millimeters) was measured in *Cybb*^f/f^ *Ncf2*^f/f^ or *Cybb Ncf2* S100A8^Cre+^ littermate controls on Day 8 post ligature in the interdental regions (M2, M2’) of the ligated molar and unligated contralateral control molars from the same mice. Data from 6-8 mice per group are displayed as mean ± SD, and statistical differences were determined using a t-test.

We also conducted a gene set enrichment analysis (GSEA) and observed that interferon signaling, G-protein-coupled receptor (GPCR) signaling, and extracellular matrix organization were positively correlated, corroborating our findings (**Figure 2E**). Neutrophils have been previously implicated in actively contributing towards inflammatory bone loss via cell-intrinsic expression of several bone resorptive ligands and also produce factors such as IL-17 that bolster the recruitment and activity of other osteoclastogenesis-driving immune cells ^28^. Several inflammatory cytokines, such as IL-1, IL-6, and TNF family members, are directly linked with osteoclastogenesis by stimulating the production of bone resorptive factors, including receptor activator of NF-κB (RANK) and its ligand (RANK-L/Tnfsf11) ^31^. Quantitative polymerase chain reaction (qPCR) analysis confirmed significantly higher expression of osteoclast-activating cytokines *Il-1b, Il-6, and Tnfsf-11* (RANK-L) in LIP gingival tissues in *Cybb^KO^,* concordant with the RNA-seq data (**Figure 2F**). Osteoprotegerin (Tnfrsf11b), which acts as a decoy ligand for RANK receptors and moderates their osteoclastic potential, was not differentially expressed between WT and *Cybb*^KO^ mice. Finally, conditionally deleting Nox2 function, specifically in neutrophils, was sufficient to drive excessive alveolar bone recession compared to littermate controls. Thus, our data indicate that the absence of Nox2 aggravates inflammatory responses at the oral mucosal barrier by dysregulating neutrophil recruitment and/or activation (**Figure 2G-H**).

### Microbial dysbiosis in *Cybb*^KO^ mice was not associated with systemic infections

Nox2 activation regulates several antimicrobial functions of neutrophils, such as phagosomal oxidative burst, degranulation, and forming neutrophil extracellular traps ^9^. We investigated whether inadequate immunosurveillance and weakened antimicrobial defense against oral bacteria drove the inflammatory burden in *Cybb*^KO^ oral tissues. The LIP model allows for profiling bacterial biofilms that accumulate around the ligatures to assess species diversity and their collective pathogenic or ‘inflammophilic’ potential in the oral cavity ^32^. 16S rRNA sequencing of ligature-associated co-housed WT and *Cybb^KO^* mice revealed that the alpha diversity Shannon Index, which measures species richness and evenness within individual samples, did not reveal significant differences between the groups. However, the beta diversity Jaccard similarity index noted a significant compositional difference between the groups (q-value = 0.001) **(Figure 3A-B).**

**Figure 3:**
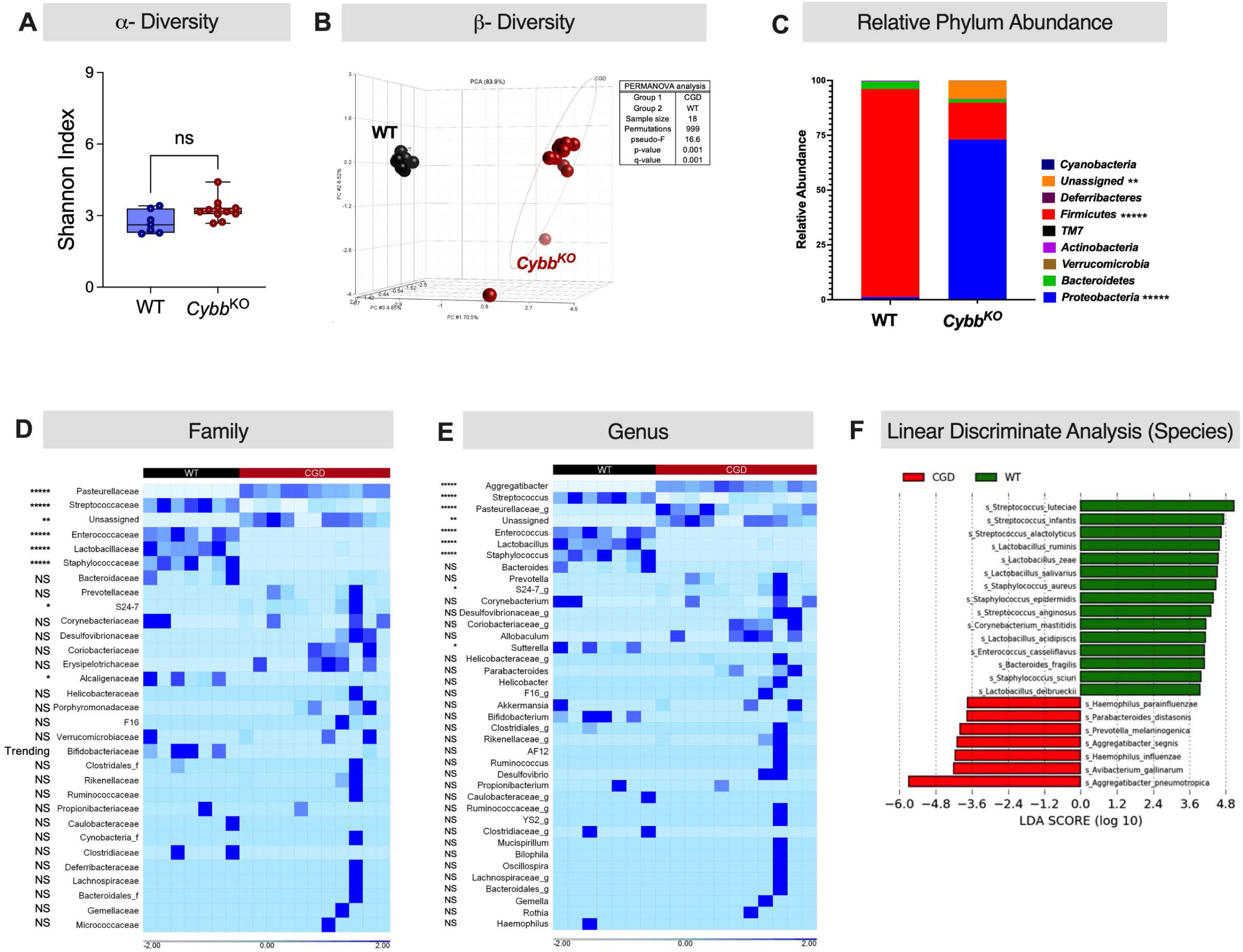
Comparative shifts in oral microbial diversity in the context of Nox2 deficiency during LIP. (**A**) Alpha diversity (species abundance and evenness) was measured with the Shannon diversity index (y-axis). Data represent mean ± SEM for n=7 for WT and n=11 for CGD mice with Mann-Whitney U test corrected p-value, ****p<0.00001. (**B**) Beta diversity (depicting compositional differences) is represented as a three-dimensional Principal Component Analysis (PCA) plot generated using Jaccard similarity matrix. PC1 vs. PC2 indicates a clear separation between *Cybb^KO^* and WT along PC1 (70.5%). PC2 vs. PC3 reveals additional variance (PC2: 8.52%, PC3: 4.85%) and clustering. PERMANOVA analysis was performed to indicate significant compositional differences (q=0.001). (**C**) Microbial composition at different taxonomic levels: (**C**) Phylum barplot (Left), (**D**) Family heatmap, and (**E**) Genus heatmap. White (light blue) signifies that microbial taxa are present in low abundance or absent, and blue signifies that microbial taxa are highly abundant. Corrected p-value derived (Dunn’s multiple testing) from Mann-Whitney U test are depicted as an asterisk next to the microbial taxa: p<0.05= *, p< 0.01 =**, p< 0.001 =***, p<0.00001=****, p<0.000001=*****, NS = not significant, and Trending = p-value between 0.051 and 0.06. **(F**) Linear Discriminate Analysis Effect size representing significant microbial species enrichment in wildtype and CGD mice. LDA cut off score >2 and p<0.05

Taxonomic analysis identified distinctive microbial communities between the groups. We found that the *Cybb^KO^* oral microbiome had increased dominance of the phylum *Proteobacteria* (typically harboring pathogens) and a concomitant reduction in the phylum *Firmicutes* (typically harboring beneficial bacteria) compared to WT mice **(Figure 3C).** Further taxonomic assessment revealed that within the phylum *Proteobacteria,* the bacterial family *Pasteurellaceae* and the genus *Aggregatibacter* were significantly expanded. At the same time, within *Firmicutes*, the genera *Streptococcus*, *Enterococcus*, *Lactobacillus*, and *Staphylococcus* in *Cybb^KO^* mice were significantly reduced **(Figure 3C).** To gain further insight into the distinguishing enrichment profiles between the two groups at the species level, linear discriminant analysis effect size (LEfSe) analyses were performed. An increase in the abundance of *Bifidobacterium pseudolongum* and other species belonging to *Firmicutes* distinguished the WT mice from *Cybb^KO^* mice. In contrast, expanding *Aggregatibacter pneumotropica*, *Allobaculum*, and *Pasteurellaceae sp.* was a hallmark feature of the microbial communities in the *Cybb^KO^* group **(Figure 3D-F).**

Although infections from *Haemophilus* and *Aggregatibacter* have been reported to cause liver abscesses and lymphadenitis in some patients with CGD, these occurrences are relatively rare ^33–35^. Typically, patients with CGD are susceptible to life-threatening infections with a narrow spectrum of pathogens. Catalase-positive bacterial pathogens (*Staphylococcus aureus*, *Burkholderia cepacia*, *Serratia marcescens*, *Nocardia* species) and specific invasive fungal pathogens (*Aspergillus* sp.) cause commonly reported CGD infections ^35^. However, recent reports indicate that although rare, specific catalase-negative pathogens such as *Chromobacterium violaceum*, *Francisella philomiragia, and Actinomyces sp* can cause significant life-threatening complications in patients with CGD ^35^. Thus, we determined whether delayed bacterial clearance contributed to oral inflammation.

Aerobic and anaerobic cultures of blood, draining lymph nodes, lungs, spleen, and liver homogenates from ligated WT and *Cybb^KO^* mice were negative, indicating no evidence of sepsis or bacterial dissemination from the oral cavity. However, given the notable differences in the oral microbiomes of WT and *Cybb^KO^* mice, we used a model of reciprocal microbiome transplant to determine relative virulence. The bacterial peritonitis model allows for quantitatively assessing antimicrobial killing capacity ^36,37^. Thus, pooled aerobic and anaerobic overnight cultures of ligature-associated bacteria were transplanted directly into the peritoneal cavities of naïve WT and *Cybb^KO^* mice as described in **Figure S2A**. While we found no significant differences in relative bacterial clearance (**Figure S2B**), *Cybb^KO^* mice had a more robust inflammatory response characterized by significantly higher neutrophil recruitment irrespective of the type of microbial challenge (**Figure S2C-G**). This data indicates that the dysregulated inflammatory responses of *Cybb^KO^* mice in the LIP model are not due to compromised antimicrobial capacity but are perhaps related to over-recruiting and/or hyperactivating ROS-deficient neutrophils.

### *Cybb^KO^* neutrophils exhibited dysregulated inflammatory responses to oral bacteria

The intensity of inflammatory responses in CGD is not always linked to active infection. In vivo challenges with bacterial ligands such as lipopolysaccharide ^38^, fungal ligands (zymosan or hyphae) ^17,18,39^, or ligands released during pyroptotic/necrotic cell death (ATP, monosodium urate crystals ^16^) have all been linked to dysregulated neutrophil recruitment and several-fold higher activation of inflammatory genes compared to WT mice. Thus, we measured neutrophil effector responses to oral bacterial stimulation using the bacterial peritonitis model, as it allows for quantitatively evaluating multiple inflammatory responses to injected bacteria (**Figure 4A**). We focused on *Porphyromonas gingivalis*, as it is a relevant pathogen in the context of human periodontal infections and is highly adept at manipulating neutrophil effector responses ^40–42^. Four hours post i.p. challenge with 10^7^ CFU of *P. gingivalis, Cybb^KO^* mice had significantly higher neutrophilic recruitment (**Figure 4C**), which correlated with significantly elevated levels of neutrophil-attracting chemokines (CXCL1, CXCL2, and LIX), as well as pro-inflammatory cytokines (IL-1α, IL-1β, TNF, and IL-12p70) in the peritoneal lavage fluid (**Figure 4B**). While we saw a significant increase in monocyte chemoattractant CCL2, peritoneal monocyte numbers 4 h post-challenge revealed no differences (**Figure 4C**). Transcriptional profiling of isolated peritoneal exudate neutrophils also suggested an increased presence of pro-inflammatory transcripts, indicating that Nox2-deficient neutrophils actively contribute towards establishing feed-forward inflammatory loops in a cell-intrinsic manner (**Figure S3A-C**). Excessive inflammatory activation was also confirmed in purified naive bone marrow neutrophils (BMNs), where ex vivo challenge with *P. gingivalis,* as well as other periodontal pathogens, led to significantly higher TNF production (**Figure S3D-F**). Interestingly, our data demonstrate more efficient clearance of *P. gingivalis* in *Cybb^KO^* peritoneal cavities, indicating that the dysregulated inflammatory response was not correlated with defective bacterial clearance (**Figure 4D**). These observations corroborate our findings with the reciprocal oral microbiome challenge (Figure S2) and are not limited to a single bacterium. In the context of *P. gingivalis,* we have previously established that Nox2 activation is not required for the intracellular killing of *P. gingivalis* by neutrophils ^40^.

**Figure 4:**
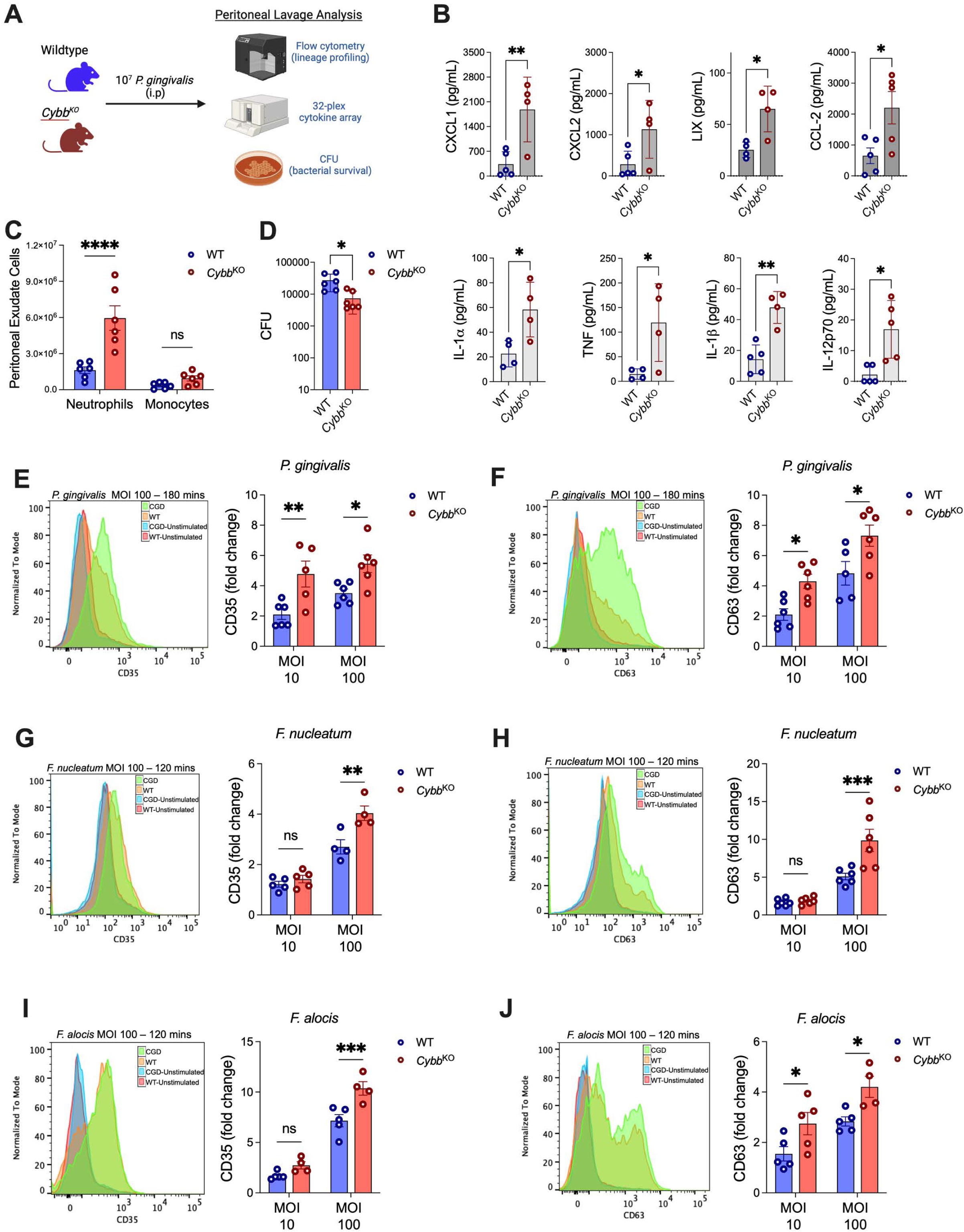
Nox2 deficiency drives excessive inflammatory cytokine production and degranulation in neutrophils. **(A)** Wild-type (WT) and *Cybb^KO^* mice were intraperitoneally challenged with 10^7^ CFU of *P. gingivalis*. After 4 h, peritoneal cavities were lavaged, and peritoneal lavage fluid was analyzed. **(B)** Cytokine and chemokine levels in peritoneal lavage fluid. (**C**) Peritoneal cell numbers in lavage fluids were identified by flow cytometry, and neutrophils (Ly6G^hi^ Ly6C^int^) and monocytes (Ly6C^hi^ Ly6G^−^). (**D**) *P. gingivalis* numbers after 4 h were determined by plating on blood agar plates, and CFUs were estimated. (**E-J**) Bone marrow neutrophils (BMNs) from WT and *Cybb^KO^*mice were challenged with oral bacteria at the indicated multiplicity of infection (MOI) for 2-3 h. Exocytosis of tertiary and azurophilic granules was quantified by measuring the surface abundance of CD35 and CD63, respectively by flow cytometry. % shifts are illustrated in histograms, and fold change from 3-7 mice (BMNs) is illustrated in adjacent bar graphs. Data from 4-6 mice per group are displayed as mean ± SD, and statistical differences were determined using a t-test or two-way ANOVA with Sidak’s multiple comparison test for grouped variables (*P<0.05; **P<0.01; ***P<0.001; ****P<0.0001).

Besides generating inflammatory cytokines, neutrophils also undergo degranulation, releasing antimicrobial mediators and proteases that can damage tissues. Excessive neutrophil degranulation has been implicated in collateral tissue damage and chronic inflammation ^43,44^. Neutrophils from patients with CGD hyper-degranulate in response to inflammatory cytokines and other agonists ^10^. Thus, we compared relative degranulation responses to oral periodontal pathogens WT and *Cybb^KO^* BMNs. As expected, *Cybb^KO^*BMNs exhibited significantly higher mobilization of primary (azurophilic) and tertiary (gelatinase) granules to the surface in response to *P. gingivalis* as well as other gram-negative (*Fusobacterium nucleatum*) and gram-positive (*Filifactor alocis*) periodontal pathogens (**Figure 4E-J**). Thus, our data demonstrate that Nox2 deficiency resulted in a broad dysregulation of neutrophil effector functions, resulting in excessive degranulation and pro-inflammatory cytokine production in response to the unbalanced microbiome, which accumulated upon placing ligatures in mice. Next, we focused on identifying transcription factors driving excessive inflammatory cytokine responses in *Cybb^KO^* mice.

### Neutrophilic oxygen consumption and Nrf2 activation are essential for limiting inflammatory damage at the oral mucosal barrier

At barrier surfaces, Nox2 activation is directly linked with activating redox-sensitive transcription factors that counter pro-inflammatory pathways and promote wound healing. For example, in the inflamed colonic mucosa, transmigrating neutrophils undergo oxidative burst, inducing “inflammatory hypoxia” and the compensatory activation of hypoxia-induced transcription factor 1 alpha (HIF-1α) in epithelial cells, which then facilitates epithelial restitution ^45^. Nox2 activation is also critical for activating another transcription factor, the Nuclear factor-erythroid 2-related factor 2 (Nrf2), a master regulator of antioxidant responses. Nuclear translocation of Nrf2 and binding to the antioxidant response element in the promoter regions drive the transcription of multiple antioxidant genes. Independently, Nrf2 upon nuclear translocation, also trans represses inflammatory transcription factors such as NF-κB ^13,38^, thereby inhibiting the transcription of IL-6 and IL-1 independent of the ARE binding motif ^46^.

In our RNA-seq data, we did not observe any differences in the HIF-1α-regulated pathways but observed suppressed Nrf2 expression (*Nfe2l2*) transcripts in the inflamed tissues of *Cybb^KO^* mice (**Figure S4A**). We further confirmed the expression *Nfe2l2* as well as Nrf2-regulated antioxidant genes (hemeoxygenase1 (*Hmox1*), NAD(P)H quinone dehydrogenase 1 (*Nqo1*), and Glutamate-cysteine ligase (*Gclc*)) by qPCR, indicating a broad suppression of Nrf2 activity in the oral tissues of *Cybb^KO^*mice (**Figure S4B**). Interestingly, our data are consistent with previous reports in which deficient Nrf2 activation in CGD mouse lungs was linked to excessive NF-κB driven hyperinflammation and pro-inflammatory cytokine expression ^13,38^. Therefore, we determined whether restoring Nrf2 activity using a synthetic agonist would ameliorate oral inflammation in *Cybb^KO^* mice.

In oral tissues, Nrf2 activation is crucial for protecting against oxidative damage, attenuating osteoclastogenesis, and promoting wound healing via the coordinated upregulation of antioxidant genes that restore cellular homeostasis ^47–49^. Unlike WT and *Cybb^KO^* mice, placing ligatures in Nrf2 knockout mice (*Nfe2l2*^−/−^) resulted in an early and severe inflammatory response, and mice had to be euthanized and removed from the study due to meeting the endpoint or humane criteria (**Figure 5A**). These observations confirmed that failing to optimally activate Nrf2 in the oral mucosa could have devastating consequences. We used sulforaphane (SFN) ^13,26^, a synthetic agonist for Nrf2, to restore Nrf2 activity and alleviate inflammatory bone loss in *Cybb^KO^* mice. SFN treatment restored the significantly diminished expression of Nrf2 and Nrf2-regulated genes *Hmox1*, *Gclc*, and *Nqo1* (**Figure 5B-E**) in the gingival tissues of inflamed *Cybb^KO^* mice and considerably reduced alveolar bone recession to levels seen in WT mice (**Figure 5F-G**).

**Figure 5:**
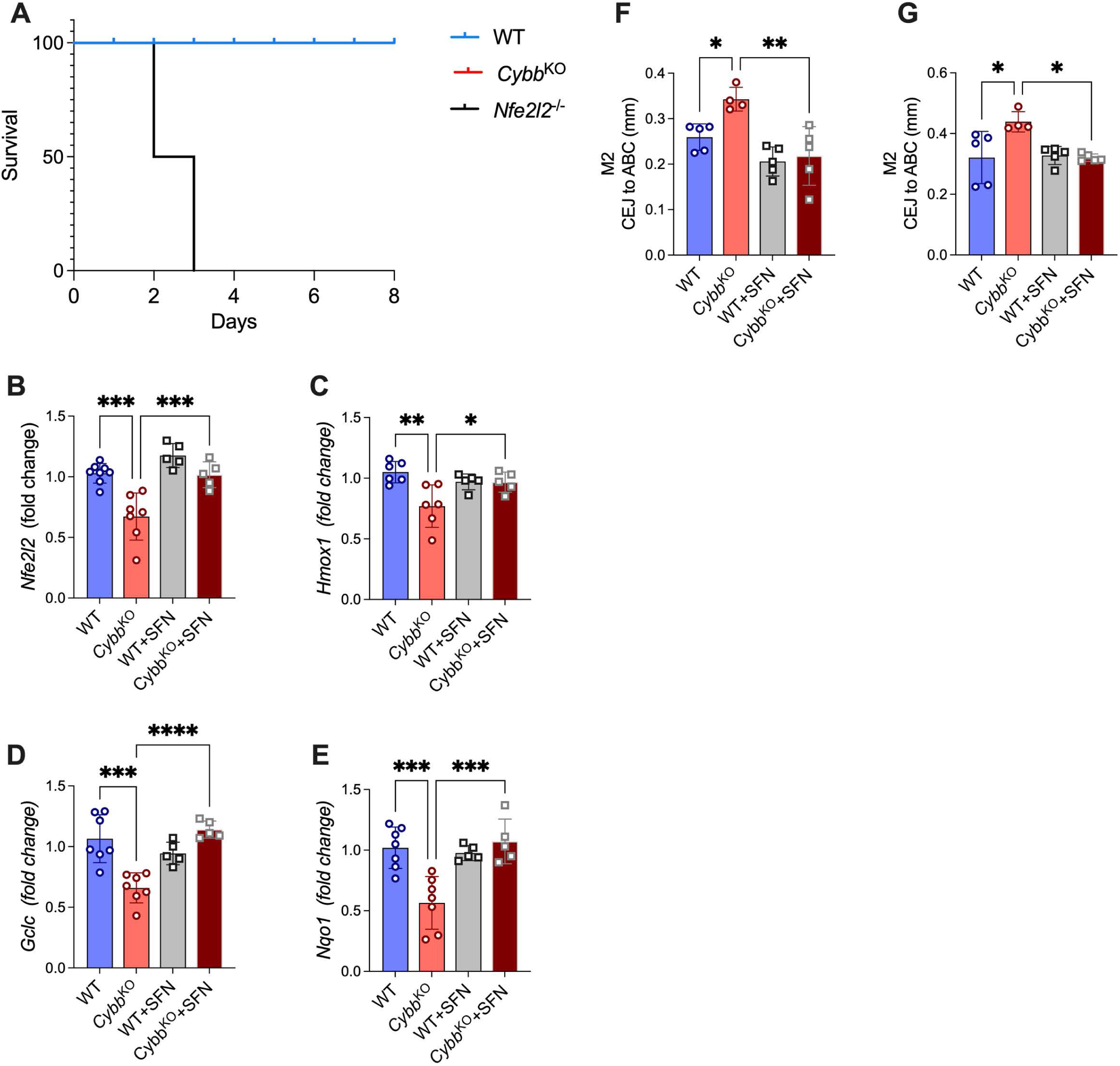
Insufficient activation of Nrf2 and downstream antioxidant responses exacerbate oral inflammation in Nox2-deficient mice. **(A)** Kaplan-Meier survival curve of wild-type (**WT**), *Cybb^KO,^* and Nrf2 knockout (*Nfe2l2*^−/−^) mice during LIP. (**B-G**) WT and *Cybb^KO^* mice were treated with 25 mg/kg sulforaphane daily for the entire duration of LIP. (**B-E**) Gingival tissues were extracted on Day 8, and **qPCR** analysis determined the relative expression of Nrf2-activated genes. (**F-G**) Average bone loss (in millimeters) in the interdental regions (M2, M2’) of the ligated molar and unligated contralateral control molars from the same mice. Data from 4-6 mice per group are displayed as mean ± SD, and statistical differences were determined using one-way ANOVA with Tukey’s multiple comparison test (*P<0.05; **P<0.01; ***P<0.001; ****P<0.0001).

## DISCUSSION

In the oral cavity, neutrophils are essential for immune surveillance, and their efferocytic clearance is critical for activating resolution or anti-inflammatory pathways necessary for maintaining homeostasis and wound healing in the oral tissues ^50^. Congenital defects compromising neutrophil recruitment, such as those observed in Leukocyte Adhesion Deficiency 1 (LAD-1) or severe congenital or cyclical neutropenias caused by neutrophil elastase (ELA2) deficiency, and Chédiak-Higashi syndrome, are all associated with severe periodontitis, often observed early in childhood ^30,51,52^. Oral lesions and ulceration are also prevalent in Papillon-Lefèvre syndrome caused by a deficiency in lysosomal exo-cysteine protease cathepsin C, involved in activating pro-enzymes of neutrophil-derived serine proteases, such as cathepsin G, elastase, and proteinase 3. Neutrophils from patients with Papillon-Lefèvre syndrome have compromised antimicrobial capacity ^51,52^. However, patients with CGD do not have abnormally low levels of neutrophils, or compromised granule repertoire, or defects in recruiting neutrophils into peripheral tissues such as the oral mucosa ^1^. Thus, the molecular mechanisms underlying oral complications in CGD are distinct from other genetic defects that impact neutrophil function.

Oral inflammation, particularly periodontitis, is an infection-driven disease where microbial virulence factors actively thwart immune clearance and perpetuate inflammation ^53,54^. Coupled with the higher incidence of infection in CGD, one would expect to find that the hyper-inflammatory responses in *Cybb^KO^* mice were linked to poor antimicrobial clearance. However, our findings challenge this notion and show that the increased susceptibility of *Cybb^KO^* mice to periodontitis was definitively attributed to dysregulation of the host inflammatory response, specifically to excessive recruitment and overactivation of ROS-deficient neutrophils. Nox2-deficient neutrophils cannot form neutrophil extracellular traps or mount an oxidative burst, thus making them inefficient at trapping and eliminating microbes ^9^. However, clinically, patients with CGD only exhibit susceptibility to a very limited spectrum of bacteria, indicating that other neutrophil antimicrobial mechanisms, such as the release of antimicrobial peptides and serine proteases, are efficient in inactivating a large variety of pathogens and also protect against endogenous colonizers or commensals ^35^. We do not observe any naturally occurring bone loss in naïve *Cybb^KO^* mice, further confirming that Nox2 deficiency does not naturally predispose to infections from oral pathobionts. Interestingly, we observed differences in the microbial composition of ligature-associated biofilm bacteria within the two strains under inflammatory conditions. It is likely that these differences arise from the enrichment of pathogenic species that produce virulence factors such as leukocidal toxins, proteases, or other virulence factors that enable them to survive the neutrophil onslaught. Many periodontal pathogens thrive in a chronic inflammatory and neutrophil-dominated sub-gingival environment due to their ability to actively incapacitate neutrophil-killing capacity. For example, *P. gingivalis* protease RgpB can degrade neutrophil azurophilic granule proteases, prolonging intracellular survival ^40,55^. In our microbiome transplant experiments, we did not observe defects in the killing of oral biofilm pathogens by Nox2-deficient neutrophils compared to WT neutrophils. Instead, we found excessive production of pro-inflammatory cytokines from neutrophils with absent ROS responses. These observations, coupled with the absence of systemic spread of oral microbes, argue against sub-optimal microbial clearance as an overwhelming driver of oral inflammation in *Cybb^KO^* mice.

A few ex vivo studies have shown excessive ROS generation by neutrophils isolated from the peripheral blood of patients with periodontitis and concluded that oxidative stress resulting from hyperactivation of Nox2 is a significant contributing factor to the pathophysiology of periodontitis ^56,57^. However, given the elevated systemic inflammatory status of patients with periodontitis, it is hard to determine whether Nox2 activity was a cause or consequence of neutrophil activation and inferred periodontal tissue destruction. Another significant confounding factor is the use of chemical inhibitors of ROS, such as N-acetylcysteine, that non-specifically inhibit multiple inflammatory pathways, including c-Jun N-terminal kinase, p38 MAP kinase, and NF-κB, thus complicating the interpretation of these studies ^58^. In contrast to these papers, our studies relied on a precise genetic deletion of Nox2 activity to demonstrate that the Nox2-derived ROS are, in fact, necessary to provide negative feedback that restrains neutrophil hyperactivation in the context of oral inflammation, damage, and dysbiosis. Our findings are aligned with several other reports that confirm the newly emerging role of Nox2 in restraining the magnitude of host inflammatory responses ^9,19,59^. Nox2 activity was essential for activating Nrf2 and Nrf2-mediated downregulation of the oral inflammatory response. Future studies will investigate whether Nrf2 restrains neutrophil-specific responses or impacts multiple cell types within oral tissues to subdue inflammation.

## AUTHOR CONTRIBUTIONS

SJ, RS, JL, KNC, KAC, and JB did all the experiments and analyzed the data. MT and MNA analyzed RNA-seq data. RB, VJ, RAI, MCD, and RJL provided either reagents, microbial strains, mice, and/or technical expertise. JB conceptualized and designed the study and wrote the manuscript with input from all authors.

## ACKNOWLEDGEMENTS

These studies were funded by DE028296 and DE028031 to JB and GM125504 to RJL.

## REFERENCES

1 Dinauer, M. C. Inflammatory consequences of inherited disorders affecting neutrophil function. Blood 133, 2130–2139 (2019). 10.1182/blood-2018-11-844563

2 Nauseef, W. M. The phagocyte NOX2 NADPH oxidase in microbial killing and cell signaling. Curr Opin Immunol 60, 130–140 (2019). 10.1016/j.coi.2019.05.006

3 Kragballe, K., Borregaard, N., Brandrup, F., Koch, C. & Staehrjohansen, K. Relation of monocyte and neutrophil oxidative metabolism to skin and oral lesions in carriers of chronic granulomatous disease. Clin Exp Immunol 43, 390–398 (1981).

4 Brandrup, F., Koch, C., Petri, M., Schiodt, M. & Johansen, K. S. Discoid lupus erythematosus-like lesions and stomatitis in female carriers of X-linked chronic granulomatous disease. Br J Dermatol 104, 495–505 (1981). 10.1111/j.1365-2133.1981.tb08163.x

5 Thomsen, I., Dulek, D. E., Creech, C. B., Graham, T. B. & Williams, J. V. Chronic granulomatous disease masquerading as Behcet disease: a case report and review of the literature. Pediatr Infect Dis J 31, 529–531 (2012). 10.1097/INF.0b013e3182481ed9

6 Dar-Odeh, N. S. et al. Orofacial findings in chronic granulomatous disease: report of twelve patients and review of the literature. BMC Res Notes 3, 37 (2010). 10.1186/1756-0500-3-37

7 Moutsopoulos, N. M. & Konkel, J. E. Tissue-Specific Immunity at the Oral Mucosal Barrier. Trends Immunol 39, 276–287 (2018). 10.1016/j.it.2017.08.005

8 Kim, T. S. & Moutsopoulos, N. M. Neutrophils and neutrophil extracellular traps in oral health and disease. Exp Mol Med 56, 1055–1065 (2024). 10.1038/s12276-024-01219-w

9 Zeng, M. Y., Miralda, I., Armstrong, C. L., Uriarte, S. M. & Bagaitkar, J. The roles of NADPH oxidase in modulating neutrophil effector responses. Mol Oral Microbiol 34, 27–38 (2019). 10.1111/omi.12252

10 Potera, R. M. et al. Neutrophil azurophilic granule exocytosis is primed by TNF-alpha and partially regulated by NADPH oxidase. Innate Immun 22, 635–646 (2016). 10.1177/1753425916668980

11 Song, Z. et al. NADPH oxidase controls pulmonary neutrophil infiltration in the response to fungal cell walls by limiting LTB4. Blood 135, 891–903 (2020). 10.1182/blood.2019003525

12 Rada, B. K. et al. Calcium signalling is altered in myeloid cells with a deficiency in NADPH oxidase activity. Clin Exp Immunol 132, 53–60 (2003). 10.1046/j.1365-2249.2003.02138.x

13 Segal, B. H. et al. NADPH oxidase limits innate immune responses in the lungs in mice. PLoS One 5, e9631 (2010). 10.1371/journal.pone.0009631

14 Yoo, D. G., Paracatu, L. C., Xu, E., Lin, X. & Dinauer, M. C. NADPH Oxidase Limits Collaborative Pattern-Recognition Receptor Signaling to Regulate Neutrophil Cytokine Production in Response to Fungal Pathogen-Associated Molecular Patterns. J Immunol 207, 923–937 (2021). 10.4049/jimmunol.2001298

15 de Luca, A. et al. IL-1 receptor blockade restores autophagy and reduces inflammation in chronic granulomatous disease in mice and in humans. Proc Natl Acad Sci U S A 111, 3526–3531 (2014). 10.1073/pnas.1322831111

16 Bagaitkar, J. et al. NADPH oxidase controls neutrophilic response to sterile inflammation in mice by regulating the IL-1alpha/G-CSF axis. Blood 126, 2724–2733 (2015). 10.1182/blood-2015-05-644773

17 Song, Z. et al. NADPH oxidase 2 limits amplification of IL-1beta-G-CSF axis and an immature neutrophil subset in murine lung inflammation. Blood Adv 7, 1225–1240 (2023). 10.1182/bloodadvances.2022007652

18 Morgenstern, D. E., Gifford, M. A., Li, L. L., Doerschuk, C. M. & Dinauer, M. C. Absence of respiratory burst in X-linked chronic granulomatous disease mice leads to abnormalities in both host defense and inflammatory response to Aspergillus fumigatus. J Exp Med 185, 207–218 (1997). 10.1084/jem.185.2.207

19 Fernandez-Boyanapalli, R. F. et al. Impaired apoptotic cell clearance in CGD due to altered macrophage programming is reversed by phosphatidylserine-dependent production of IL-4. Blood 113, 2047–2055 (2009). 10.1182/blood-2008-05-160564

20 Petersen, J. E. et al. Enhanced cutaneous inflammatory reactions to Aspergillus fumigatus in a murine model of chronic granulomatous disease. J Invest Dermatol 118, 424–429 (2002). 10.1046/j.0022-202x.2001.01691.x

21 Pollock, J. D. et al. Mouse model of X-linked chronic granulomatous disease, an inherited defect in phagocyte superoxide production. Nat Genet 9, 202–209 (1995). 10.1038/ng0295-202

22 Idol, R. A. et al. Neutrophil and Macrophage NADPH Oxidase 2 Differentially Control Responses to Inflammation and to Aspergillus fumigatus in Mice. J Immunol (2022). 10.4049/jimmunol.2200543

23 Abe, T. & Hajishengallis, G. Optimization of the ligature-induced periodontitis model in mice. J Immunol Methods 394, 49–54 (2013). 10.1016/j.jim.2013.05.002

24 Dutzan, N., Abusleme, L., Konkel, J. E. & Moutsopoulos, N. M. Isolation, Characterization and Functional Examination of the Gingival Immune Cell Network. J Vis Exp, 53736 (2016). 10.3791/53736

25 Park, C. H. et al. Three-dimensional micro-computed tomographic imaging of alveolar bone in experimental bone loss or repair. J Periodontol 78, 273–281 (2007). 10.1902/jop.2007.060252

26 Ma, C. et al. Sulforaphane alleviates psoriasis by enhancing antioxidant defense through KEAP1-NRF2 Pathway activation and attenuating inflammatory signaling. Cell Death Dis 14, 768 (2023). 10.1038/s41419-023-06234-9

27 Furze, R. C. & Rankin, S. M. Neutrophil mobilization and clearance in the bone marrow. Immunology 125, 281–288 (2008). 10.1111/j.1365-2567.2008.02950.x

28 Hajishengallis, G., Moutsopoulos, N. M., Hajishengallis, E. & Chavakis, T. Immune and regulatory functions of neutrophils in inflammatory bone loss. Semin Immunol 28, 146–158 (2016). 10.1016/j.smim.2016.02.002

29 Hajishengallis, G. Illuminating the oral microbiome and its host interactions: animal models of disease. FEMS Microbiol Rev 47 (2023). 10.1093/femsre/fuad018

30 Jung, S., Gies, V., Korganow, A. S. & Guffroy, A. Primary Immunodeficiencies With Defects in Innate Immunity: Focus on Orofacial Manifestations. Front Immunol 11, 1065 (2020). 10.3389/fimmu.2020.01065

31 Redlich, K. & Smolen, J. S. Inflammatory bone loss: pathogenesis and therapeutic intervention. Nat Rev Drug Discov 11, 234–250 (2012). 10.1038/nrd3669

32 Hajishengallis, G. The inflammophilic character of the periodontitis-associated microbiota. Mol Oral Microbiol 29, 248–257 (2014). 10.1111/omi.12065

33 Kottilil, S., Malech, H. L., Gill, V. J. & Holland, S. M. Infections with Haemophilus species in chronic granulomatous disease: insights into the interaction of bacterial catalase and H2O2 production. Clin Immunol 106, 226–230 (2003). 10.1016/s1521-6616(02)00048-7

34 Dalby, S., Andersen, T. L., Greisen, P. W., Petersen, H. & Husby, S. Abdominal Positron Emission Tomography Combined With Magnetic Resonance Imaging in Chronic Granulomatous Disease. JPGN Rep 2, e047 (2021). 10.1097/PG9.0000000000000047

35 Yu, J. E., Azar, A. E., Chong, H. J., Jongco, A. M., 3rd & Prince, B. T. Considerations in the Diagnosis of Chronic Granulomatous Disease. J Pediatric Infect Dis Soc 7, S6–S11 (2018). 10.1093/jpids/piy007

36 Sandberg, A., Hessler, J. H., Skov, R. L., Blom, J. & Frimodt-Moller, N. Intracellular activity of antibiotics against Staphylococcus aureus in a mouse peritonitis model. Antimicrob Agents Chemother 53, 1874–1883 (2009). 10.1128/AAC.01605-07

37 Knudsen, J. D., Frimodt-Moller, N. & Espersen, F. Experimental Streptococcus pneumoniae infection in mice for studying correlation of in vitro and in vivo activities of penicillin against pneumococci with various susceptibilities to penicillin. Antimicrob Agents Chemother 39, 1253–1258 (1995). 10.1128/AAC.39.6.1253

38 Han, W. et al. NADPH oxidase limits lipopolysaccharide-induced lung inflammation and injury in mice through reduction-oxidation regulation of NF-kappaB activity. J Immunol 190, 4786–4794 (2013). 10.4049/jimmunol.1201809

39 Fernandez-Boyanapalli, R. et al. PPARgamma activation normalizes resolution of acute sterile inflammation in murine chronic granulomatous disease. Blood 116, 4512–4522 (2010). 10.1182/blood-2010-02-272005

40 Cooper, K. N. et al. Limited proteolysis of neutrophil granule proteins by the bacterial protease RgpB depletes neutrophil antimicrobial capacity. J Leukoc Biol (2024). 10.1093/jleuko/qiae209

41 Bryzek, D. et al. Triggering NETosis via protease-activated receptor (PAR)-2 signaling as a mechanism of hijacking neutrophils function for pathogen benefits. PLoS Pathog 15, e1007773 (2019). 10.1371/journal.ppat.1007773

42 Sochalska, M. & Potempa, J. Manipulation of Neutrophils by Porphyromonas gingivalis in the Development of Periodontitis. Front Cell Infect Microbiol 7, 197 (2017). 10.3389/fcimb.2017.00197

43 Othman, A., Sekheri, M. & Filep, J. G. Roles of neutrophil granule proteins in orchestrating inflammation and immunity. FEBS J 289, 3932–3953 (2022). 10.1111/febs.15803

44 Liew, P. X. & Kubes, P. The Neutrophil’s Role During Health and Disease. Physiol Rev 99, 1223–1248 (2019). 10.1152/physrev.00012.2018

45 Campbell, E. L. et al. Transmigrating neutrophils shape the mucosal microenvironment through localized oxygen depletion to influence resolution of inflammation. Immunity 40, 66–77 (2014). 10.1016/j.immuni.2013.11.020

46 Kobayashi, E. H. et al. Nrf2 suppresses macrophage inflammatory response by blocking proinflammatory cytokine transcription. Nat Commun 7, 11624 (2016). 10.1038/ncomms11624

47 Kataoka, K. et al. Visualization of Oxidative Stress Induced by Experimental Periodontitis in Keap1-Dependent Oxidative Stress Detector-Luciferase Mice. Int J Mol Sci 17 (2016). 10.3390/ijms17111907

48 Kanzaki, H. et al. Nuclear Nrf2 induction by protein transduction attenuates osteoclastogenesis. Free Radic Biol Med 77, 239–248 (2014). 10.1016/j.freeradbiomed.2014.09.006

49 Sima, C. et al. Nuclear Factor Erythroid 2-Related Factor 2 Down-Regulation in Oral Neutrophils Is Associated with Periodontal Oxidative Damage and Severe Chronic Periodontitis. Am J Pathol 186, 1417–1426 (2016). 10.1016/j.ajpath.2016.01.013

50 Silva, L. M., Kim, T. S. & Moutsopoulos, N. M. Neutrophils are gatekeepers of mucosal immunity. Immunol Rev 314, 125–141 (2023). 10.1111/imr.13171

51 Hajishengallis, G., Chavakis, T., Hajishengallis, E. & Lambris, J. D. Neutrophil homeostasis and inflammation: novel paradigms from studying periodontitis. J Leukoc Biol 98, 539–548 (2015). 10.1189/jlb.3VMR1014-468R

52 de Arruda, J. A. A. et al. Oral manifestations of Chediak-Higashi syndrome: A systematic review. Dis Mon 69, 101356 (2023). 10.1016/j.disamonth.2022.101356

53 Lamont, R. J. & Hajishengallis, G. Polymicrobial synergy and dysbiosis in inflammatory disease. Trends Mol Med 21, 172–183 (2015). 10.1016/j.molmed.2014.11.004

54 Lamont, R. J., Koo, H. & Hajishengallis, G. The oral microbiota: dynamic communities and host interactions. Nat Rev Microbiol 16, 745–759 (2018). 10.1038/s41579-018-0089-x

55 Cooper, K. N. et al. The trapping of live neutrophils by macrophages during infection. Cell Death Dis 16, 488 (2025). 10.1038/s41419-025-07808-5

56 Chapple, I. L. & Matthews, J. B. The role of reactive oxygen and antioxidant species in periodontal tissue destruction. Periodontol 2000 43, 160–232 (2007). 10.1111/j.1600-0757.2006.00178.x

57 Matthews, J. B., Wright, H. J., Roberts, A., Cooper, P. R. & Chapple, I. L. Hyperactivity and reactivity of peripheral blood neutrophils in chronic periodontitis. Clin Exp Immunol 147, 255–264 (2007). 10.1111/j.1365-2249.2006.03276.x

58 Zafarullah, M., Li, W. Q., Sylvester, J. & Ahmad, M. Molecular mechanisms of N-acetylcysteine actions. Cell Mol Life Sci 60, 6–20 (2003). 10.1007/s000180300001

59 Bagaitkar, J. et al. NADPH oxidase activation regulates apoptotic neutrophil clearance by murine macrophages. Blood 131, 2367–2378 (2018). 10.1182/blood-2017-09-809004

