## Supplemental methods and figures for "The Nox2 NADPH oxidase regulates neutrophilic inflammation in the oral cavity"

_Isolation and purification of bone marrow neutrophils_: Bone marrow cells were obtained by flushing the tibias and femurs of mice, and bone marrow neutrophils (BMNs) were purified using the anti-Ly6G microbead kit by positive selection per the manufacturer's protocol (Miltenyi). Cytospins and Wright Giemsa staining assessed percent purity.

_ELISAs and Cytokine arrays_: Cell-free supernatants were analyzed for TNF levels using commercially available mouse TNF ELISA kits (eBiosciences). Cytokine levels in mouse serum and peritoneal lavage fluid were also assessed using Milliplex mouse cytokine/chemokine 32-plex array beads (Millipore-Sigma) and analyzed using Bio-Rad Bioplex-200 analyzer.

_Flow cytometry_: Blood (~200 µl) was isolated into EDTA vacutainers from submandibular bleeds or cardiac stick post-euthanasia. Red blood cells (RBCs) were lysed using 1x pharm lyse buffer for 3 mins. 'Gingival blocks' were dissected from mice as previously described by Dutzan et al. [26] and enzymatically digested (5 ml RPMI containing 3.2 mg/ml Collagenase VI and 0.15 µg/ml DNAse) for 30-45 mins at 37°C with gentle shaking. In the last 5 min of incubation, 50 µl of 0.5 M EDTA was used. Gingival tissues and media were passed through a 70 µ cell strainer to obtain single-cell suspensions. Peritoneal cells were isolated using PBS with 2 mM EDTA. For flow staining, all cell pellets were suspended in 100 µl Fc block (2.4 G2 supernatants) for 10 min on ice prior to the addition of antibodies. Cells were stained using a combination of rat anti-mouse antibodies: CD45-BV510 (clone 30-F11; BD Horizon), Ly6G-V450 (clone 1A8; BD Horizon), Ly6C-PerCP/Cy5.5 (clone HK1.4; Biolegend), and F4/80-APC-Cy7 (clone BM8; Biolegend). See gating strategies for subset identification. Data was collected on BD Celesta and BD Fortessa flow cytometers and analyzed using FlowJo software.

*qPCR*: Total RNA was extracted from tissues or cells using the RNeasy Plus Mini Kit (Qiagen) per the manufacturer's protocol. cDNA was synthesized by High-Capacity cDNA Reverse Transcription Kit (Applied Biosystems). qPCR was performed using TaqMan Fast Universal PCR Master Mix (ThermoFisher) on a 7500 Fast Real-Time PCR system (Applied Biosystems) using primers and Taqman probes purchased from Thermo Fisher: *Il1b* Mm00434228_m1; *Il6* Mm00446190_m1; *Tnf* Mm00443258_m1; *Tnfsf11* Mm00441906_m1; *Csf1* Mm00432686_m1; *Nfe2l2* Mm00477784_m1; *Hmox1* Mm00516005_m1; *Nqo1* Mm01253561_m1; *Gclc* Mm00802655_m1.

*16S rRNA Gene Sequencing and Metagenomic analysis:* Genomic DNA was extracted from ligatures placed in mice for 8 days using ReliaPrep™ gDNA Tissue Miniprep System (Promega Cat # A2051). A total of 10 ng of input DNA was used for PCR amplification of bacterial full-length 16S rRNA genes (V1—V9 regions) using KAPA HiFi HotStart ReadyMix PCR kit (Roche KK2600), Forward primer: 5'GCATC/barcode/AGRGTTYGATYMTGGCTCAG3' and Reverse primer: 5'GCATC/barcode/RGYTACCTTGTTACGACTT3'. PCR settings were: Initial denaturation at 95°C for 3 minutes followed by 25 cycles of denaturation at 95°C for 30 seconds, annealing at 57°C for 30 seconds, and extension at 72°C for 60 seconds. Samples were then pooled, and multiplexed amplicon libraries were prepared using SMRTbell prep kit 3 (PacBio 102-182-700) and sequenced on the PacBio RS sequel II platform. The average sequencing depth obtained was 6500 reads per sample.

The microbial diversity analysis and taxonomic profiling were done using Quantitative Insights into the Microbial Ecology (QIIME) version 2-2022.2 [1]. Demultiplexed HiFi reads were imported to qiime2-2022.2, and denoising of HiFi reads into amplicon sequence variants (ASVs) was performed using DADA2 qiime2-2022.2 plugin [2]. The taxonomic classification of ASVs was done using VSEARCH at a similarity cutoff value of 97% followed by mapping of ASVs using

Greengenes 13_8_99 classifier [3]. A phylogenetic tree was constructed using qiime phylogeny align-to-tree-mafft-fasttree algorithm. Further, an even sequence sampling depth was utilized when generating all diversity measures. Alpha diversity (diversity in individual samples) was determined using the Shannon index [4]. Beta diversity (diversity between samples) was determined using the Jaccard similarity distance matrix, and principal component analysis plots were generated using Partek genomics suite version 7.21.1119, and PERMANOVA analysis was performed to evaluate statistical differences. For the taxonomic data ASV values were converted to relative abundance using customized R script (version 4.0) and represented as bar plots and heatmaps using Partek genomics suite version 7.21.1119. Mann-Whitney U statistical tests were performed using Partek genomics suite version 7.21.1119 for taxonomic data and GraphPad PRISM version 9.4.0 for alpha diversity. Corrected p-value (<0.05) significance was obtained after Dunn's multiple corrections. Further, linear discriminant analysis Effect Size (LEfSe) was used on species relative abundance data to probe for significant species enrichment that distinguish the two groups (wildtype and CGD/ *Cybb*$^{KO}$ mice) [5]. The analysis was performed using http://huttenhower.sph.harvard.edu/galaxy. Significant species enrichment was calculated via a series of Kruskal Wallis non-parametric tests to determine significant alterations in taxonomic abundance between groups. This was followed up using linear discriminant analysis to estimate the effect size of any significant alterations set at LDA cut-off score >2 and p<0.05

*mRNA sequencing*

The TruSeq Stranded mRNA Library Prep Kit (Illumina Cat# 20020594) and TruSeq RNA CD Index Plate (Illumina Cat# 20019792) were used to generate a sequencing library from 500ng of RNA. Size, purity, and semi-quantitation were performed on an Agilent Bioanalyzer 2100 system using the Agilent DNA HS Kit. Paired-end sequencing was performed on an Illumina NextSeq 500, using the NextSeq 500/550 75 cycles High Output Kit v2.5 (Illumina Cat# 20024906) at the

University of Louisville core. Single-end reads were aligned with STAR v2.6 [6]. Each sample was evaluated according to post-alignment quality control measures with RSeQC tools [7]. Gene counts were quantified using featureCounts v2.0.1 [8] with reverse strand specificity parameter. Alignment and gene counts were generated against the GRCm38 (Ensembl release 93) genome assembly. The DESeq2 computational pipeline (version 1.34.0) [9] was used to normalize counts and perform differential expression analysis. Gene set enrichment analysis was performed on the pre-ranked list using the R package *fgsea* [10] with Canonical pathways collection for Mus Musculus from the Molecular Signatures Database, accessed with R package *msigdbr* [11]. An adjusted $p$-value < 0.05 was used as the significance threshold for the selection of significantly enriched pathways. Visualization was produced using the package *ggplot2*, and heatmaps were generated with the *pheatmap* package. Functions were implemented within the R programming environment version 4.1.2.

**SUPPLEMENTAL FIGURES:**

### Supplemental Figure S1

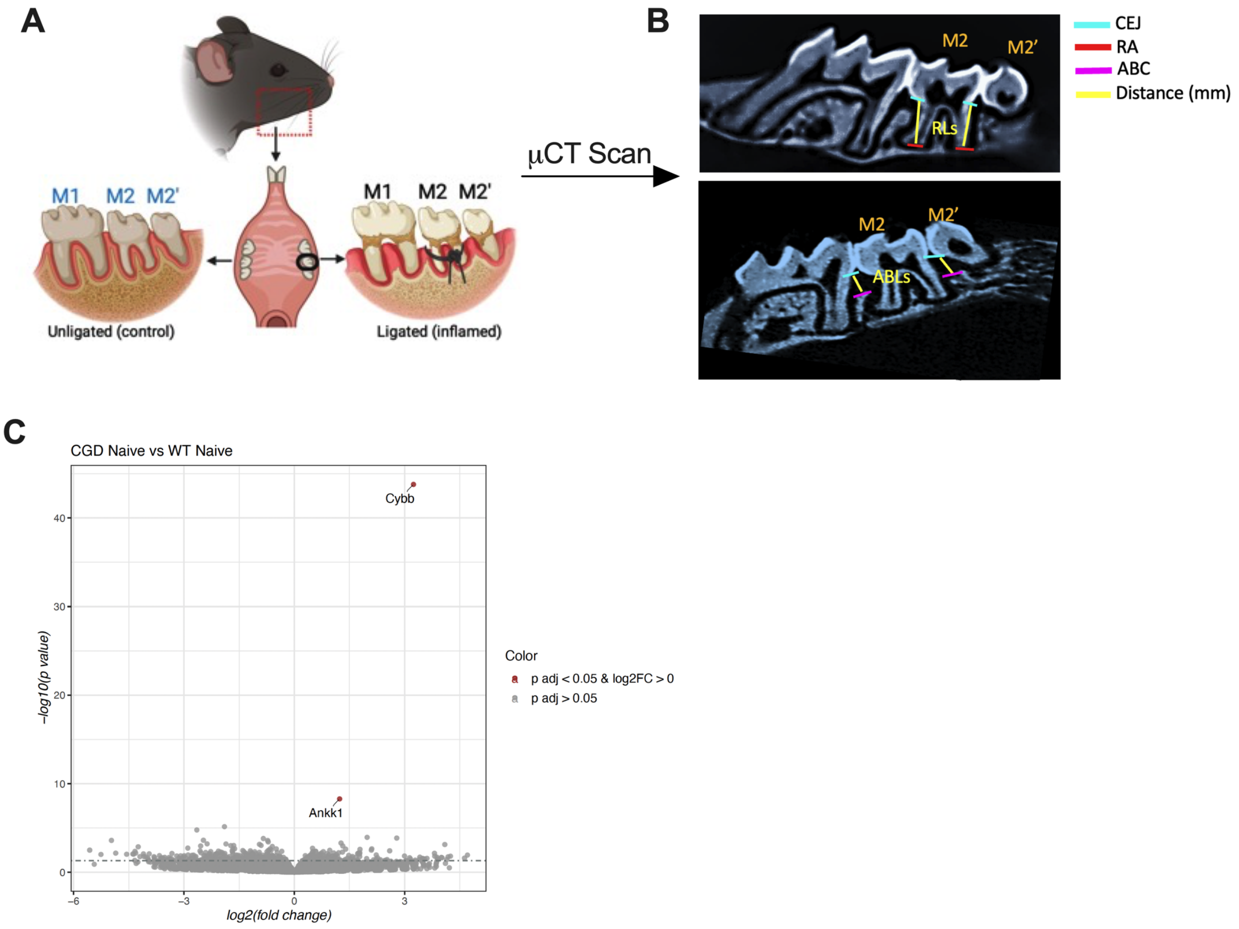


**Supplemental Figure S1:** (A, B) Illustration showing ligature placement around second molars of the maxilla. Average bone loss was measured by taking linear measurements (in millimeters) from the cemento-enamel junction (CEJ) to the alveolar bone crest (ABC) in the interdental regions between the first and second molars (M1-M2) or the second and third molars (M2-M2’). (C) Volcano plots of RNA-seq data were generated by plotting log2 (fold change) versus – log10 (adjusted P-value), representing differences between naïve WT and *Cybb*$^{KO}$ gingival tissues.

### Supplemental Figure S2

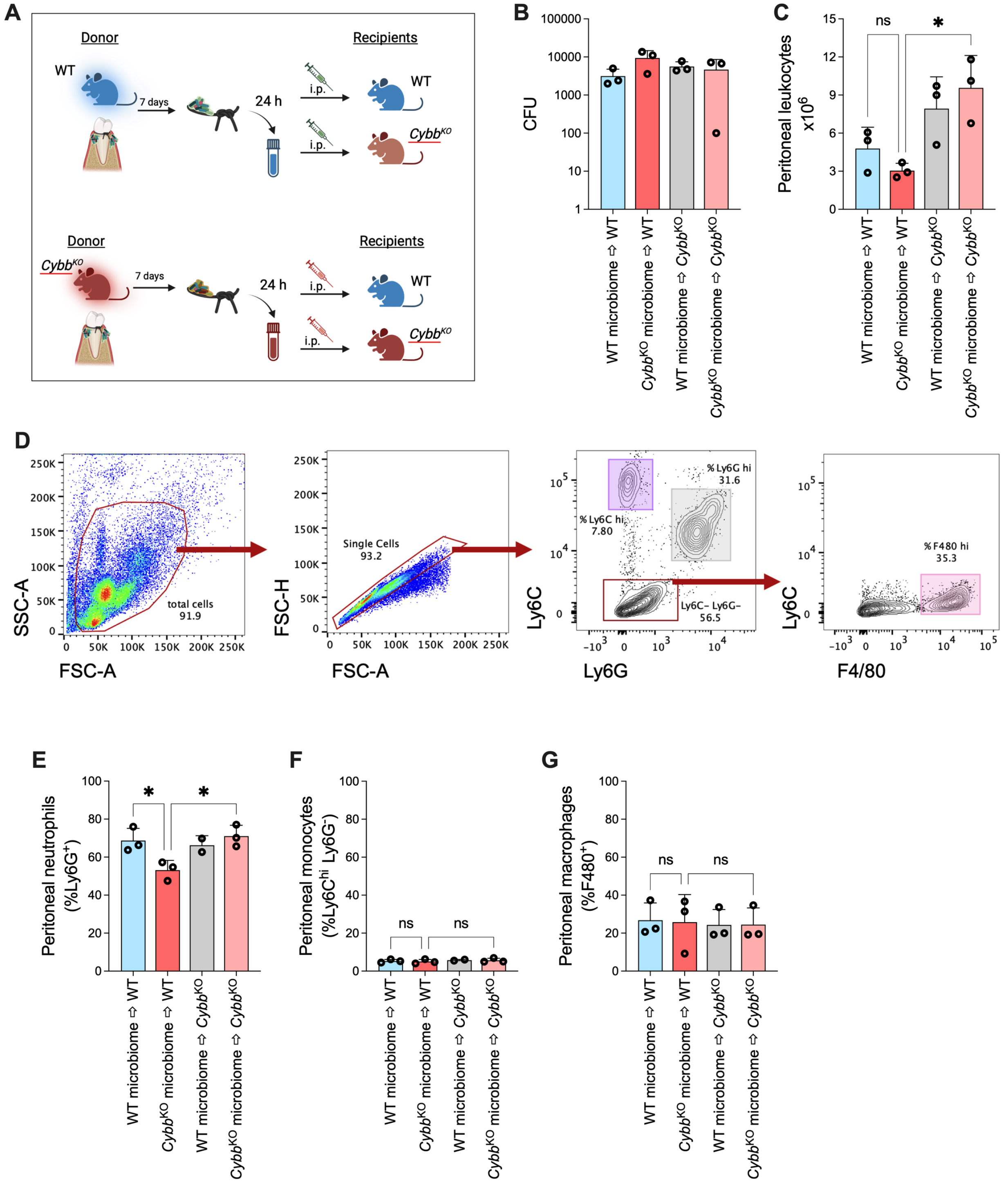


**Supplemental Figure S2:** The ligature-associated microbiome was isolated on Day 8 from WT and *Cybb*$^{KO}$ mice 'donor' mice and cultured overnight under aerobic and anaerobic conditions.

Next day, $10^7$ pooled CFUs from aerobic and anaerobic overnight cultures were injected intraperitoneally in naïve or ‘recipient’ WT and *Cybb*$^{KO}$ mice. After 4 h, peritoneal cavities were lavaged with sterile saline, and (B) peritoneal bacterial numbers (CFUs) were determined by plating on agar plates. (C)Total peritoneal cell numbers recovered. (D) Gating strategy for identification of peritoneal cells based on lineage markers. (E-G) Relative % of neutrophils (Ly6G$^{hi}$ Ly6C$^{int}$); monocytes (Ly6C$^{hi}$ Ly6G$^{-}$); and macrophages (F4/80$^{+}$) in total peritoneal cells. Data from 3 mice per group are shown as mean ± SD, and statistical differences were determined using one-way ANOVA with Tukey’s multiple comparison test (*P<0.05; **P<0.01; ***P<0.001; ****P<0.0001).

### Supplemental Figure S3

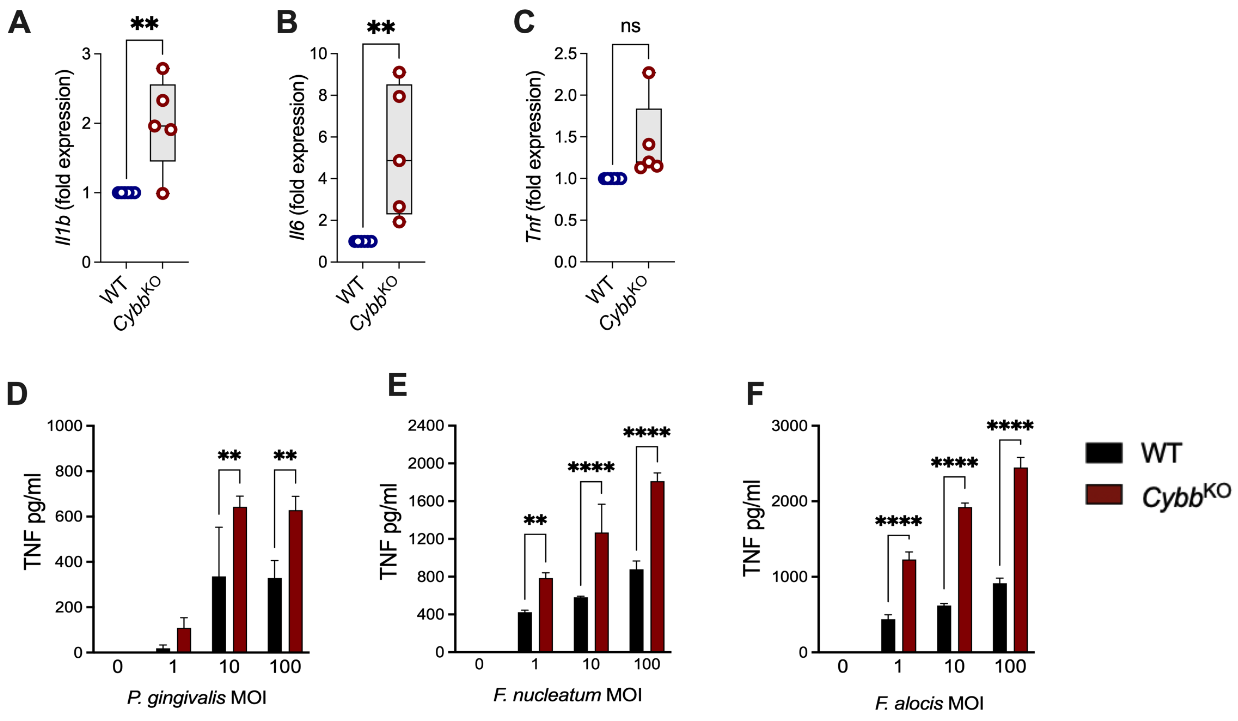


**Supplemental Figure S3:** Wildtype (WT) and *Cybb*$^{KO}$ mice were intraperitoneally challenged with $10^7$ CFU of *P. gingivalis*. After 4 h, peritoneal cavities were lavaged to isolate peritoneal neutrophils. Relative transcript expression of (A) *Il1b*, (B) *Il6,* and (C) *Tnf* was determined by qPCR in peritoneal neutrophils. Data from 5 mice per group are shown as mean ± SD, and statistical differences were determined using a t-test (*P<0.05; **P<0.01; ***P<0.001; ****P<0.0001). Bone marrow neutrophils isolated from WT and *Cybb*$^{KO}$ mice were challenged with oral bacteria at indicated MOIs for 18 h. TNF levels in cell-free supernatants were determined by ELISA. Statistical differences were determined using a two-way ANOVA with Sidak's multiple comparison test for grouped variables (*P<0.05; **P<0.01; ***P<0.001; ****P<0.0001).

### Supplemental Figure S4

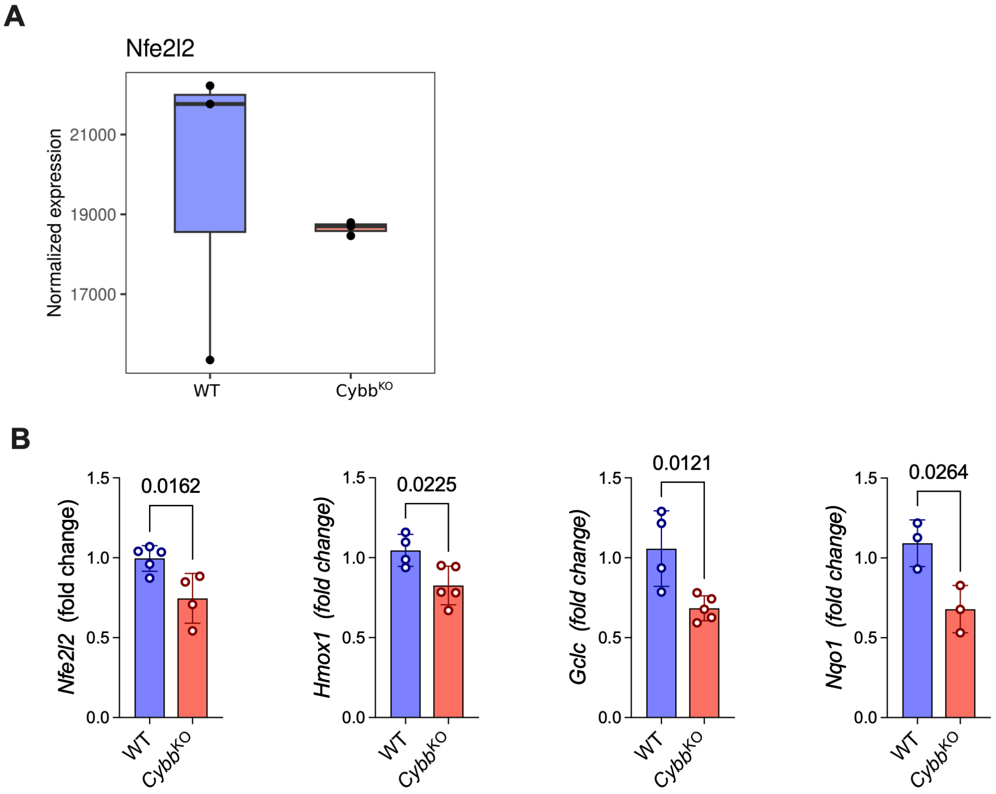


**Supplemental**

**Figure S4:** Gene expression was measured on Day 8 after ligature placement in the gingival tissues of wildtype and *Cybb*$^{KO}$ mice by RNA-seq (A) or qPCR (B). Statistical differences were determined by t-test.

**REFERENCES:**


1. Hall M, Beiko RG. 16S rRNA Gene Analysis with QIIME2. *Methods Mol Biol*. 2018;1849:113-129.

2. Callahan BJ, McMurdie PJ, Rosen MJ, Han AW, Johnson AJ, Holmes SP. DADA2: High-resolution sample inference from Illumina amplicon data. *Nat Methods*. 2016;13(7):581-583.

3. DeSantis TZ, Hugenholtz P, Larsen N, et al. Greengenes, a chimera-checked 16S rRNA gene database and workbench compatible with ARB. *Appl Environ Microbiol*. 2006;72(7):5069-5072.

4. Willis AD. Rarefaction, Alpha Diversity, and Statistics. *Front Microbiol*. 2019;10:2407.

5. Segata N, Izard J, Waldron L, et al. Metagenomic biomarker discovery and explanation. *Genome Biol*. 2011;12(6):R60.

6. Dobin A, Davis CA, Schlesinger F, et al. STAR: ultrafast universal RNA-seq aligner. *Bioinformatics*. 2013;29(1):15-21.

7. Wang L, Wang S, Li W. RSeQC: quality control of RNA-seq experiments. *Bioinformatics*. 2012;28(16):2184-2185.

8. Liao Y, Smyth GK, Shi W. featureCounts: an efficient general purpose program for assigning sequence reads to genomic features. *Bioinformatics*. 2014;30(7):923-930.

9. Love MI, Huber W, Anders S. Moderated estimation of fold change and dispersion for RNA-seq data with DESeq2. *Genome Biol*. 2014;15(12):550.

10. Korotkevich G SV, Budin N, Shpak B, Artyomov MN, Sergushichev A. Fast gene set enrichment analysis. *bioRxiv*. 2019.

11. Liberzon A, Subramanian A, Pinchback R, Thorvaldsdottir H, Tamayo P, Mesirov JP. Molecular signatures database (MSigDB) 3.0. *Bioinformatics*. 2011;27(12):1739-1740.
